# Propagation of seminal toxins through binary expression gene drives can suppress polyandrous populations

**DOI:** 10.1101/2021.11.23.469777

**Authors:** Juan Hurtado, Santiago Revale, Luciano M Matzkin

## Abstract

Gene drives can be highly effective in controlling a target population by disrupting a female fertility gene. To spread across a population, these drives require that disrupted alleles be largely recessive so as not to impose too high of a fitness penalty. We argue that this restriction may be relaxed by using a double gene drive design to spread a split binary expression system. One drive carries a dominant lethal/toxic effector alone and the other a transactivator factor, without which the effector will not act. Only after the drives reach sufficiently high frequencies would individuals have the chance to inherit both system components and the effector be expressed. We explore through mathematical modeling the potential of this design to spread dominant lethal/toxic alleles and suppress populations. We show that this system could be implemented to spread engineered seminal proteins designed to kill females, making it highly effective against polyandrous populations.

## Introduction

There are, in most parts of the world, insect species that inflict significant harm to the human condition. These insect species, usually invasive, cause or transmit diseases, reduce agricultural production, or threaten endemic biodiversity (reviewed in ^1^). However, only a few of these species can be effectively controlled by current methods, in many cases by using unsustainable or ecologically damaging strategies such as broad-spectrum pesticides. More recently, integrative pest management (IPM) has focused on minimizing the use of pesticides, while promoting other less ecologically damaging strategies, such as biological control ^2^. The great efficiency and versatility of CRISPR-Cas9 nucleases for genome editing and the super-Mendelian inheritance of self-propagating gene drives have recently made engineered CRISPR-Cas9-mediated (gene) drives one of the most promising—though controversial—technologies for biological control ^3–10^. This young technology could be used, for instance, to quickly spread desired genetics variants (e.g., payload genes) through a target population after a single release of transgenic individuals carrying the engineered drive.

During meiosis in sexually reproducing species, heterozygous individuals containing a single drive allele can have it copied onto the homolog wild-type (wt) chromosome in a process known as drive conversion or homing ^3^. Homing can increase the inheritance rate of gene drive alleles beyond the Mendelian rule of gene segregation, pushing it towards 100%. Therefore, a drive can spread through a population even if it confers a fitness cost ^3^. Based on this mechanism, fertility or viability suppression drives are intended to introduce a considerable genetic load into the target population by homing and disrupting a gene essential for fertility or viability. The potential of this approach has been shown in silico through mathematical modeling (e.g., ^11–17^). A suppression drive targeting the *doublesex* gene, which controls sex determination and is essential for female fertility in mosquitoes ^18^, is among the most effective gene drive systems recently developed in the laboratory (e.g., ^7,10,19,20^). This *doublesex* gene drive was able to collapse cage populations of the *Anopheles gambiae* mosquito in as few as seven generations ^18^. However, a number of technical limitations still call into question the versatility of this approach in the field ^21–28^.

One of the biggest limitations to the spread of a single gene drive system (affecting fertility or viability) is the requirement that the disrupted allele effects be largely recessive and preferentially target only females, i.e., the target gene should be haplosufficient for females while, for the best outcome, nonessential for males ^12,15^. If the drive carries a dominant or semidominant lethal gene, which may avoid the need of targeting haplosufficient endogenous genes, it imposes a high fitness penalty and, consequently, it may not be able to invade the population. In many species, however, finding haplosufficient endogenous genes may be challenging, not only because it requires a deep and comprehensive knowledge of female and male fertility/viability genetics but also because many of the essential genes involved in female fertility/viability may affect pleiotropically other life-history traits and/or may not be largely haplosufficient (i.e., mutant heterozygotes may experience reduced fertility/viability), both of which impede drive spread.

We propose that the haplosufficiency limitation of suppression drives may be significantly reduced by deploying a double gene drive design to spread a split binary expression system. One drive would carry a transactivator factor (Fig. 1A) and the other would carry a dominant or semidominant lethal/toxic effector (Fig. 1B). As part of a binary expression system, the effector would only be expressed in the presence of the transactivator protein, so that such system would impose a low fitness cost at its introduction and may allow each drive alone to rapidly spread from rare. After some generations, only when each drive reaches sufficiently high frequencies, both system components would have the chance to co-occur within an individual. At this point the effector would be expressed killing or sterilizing the individual, eventually leading to a decline of the target population. We refer to this double gene drive design as the Binary Expression Drive (BED). The concept is similar to the ‘interacting drives’ approach proposed by Esvelt and co-workers ^6^. However, while the ‘interacting drives’ approach achieves suppression effects by disrupting native, essential genes, BED aims to cause the expression of non-native, dominant toxins.

**Fig. 1.**
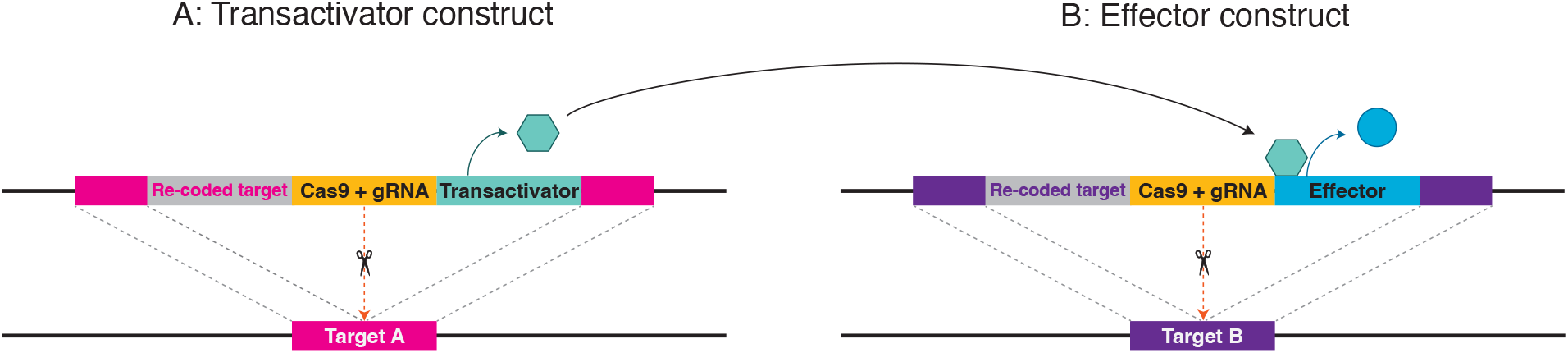
Binary Expression Drive constructs. Left: the transactivator construct (A). Right: the effector construct (B). The promoter of the transactivator gene is only active in the target tissue/organ, for instance, the ovaries or the male accessory glands. The transactivator protein will activate the expression of the effector gene, which encodes a dominant or semidominant toxin, in the target tissue/organ. In both constructs, the expression of Cas9 and the gRNA is under the regulation of a germline promoter that allows drive conversion during meiosis. Ideally, target loci are highly conserved genes that cannot tolerate mutations so resistance alleles are selected out of the population while each drive allele alone, carrying a re-coded target gene, will not impose an intended fitness cost.

BED can be used, for instance, to spread intracellular toxic proteins to be specifically expressed in the female reproductive organs so adult females carrying both the transactivator and the effector constructs would be sterilized. We will refer to this design as ‘female fertility BED’ (ffBED). In polyandrous species, where females mate with several males, the BED concept could be modified to spread engineered seminal proteins designed to be activated in the uterus and kill or sterilize females. We will refer to this design as a ‘seminal BED’ (sBED). Males carrying a sBED will behave as “terminators” of females, transferring into them lethal toxins.

Here, we developed a model for stochastic simulations in R language to study the behavior of BEDs and compare it to female fertility single drives (ffSD) targeting a haplosufficient but essential female fertility gene. In both BED designs described here (female fertility or seminal), the transactivator and effector constructs are inserted into unlinked autosomal loci that are essential for embryonic development. In the ffBEDs, the transactivator construct is exclusively expressed in the ovary, and the effector construct encodes a toxin that sterilizes the female. In the sBEDs, the transactivator construct is only expressed in the male accessory gland, and the effector construct encodes a seminal toxin that is innocuous to the male that produces it but lethal to the female that receives it via the ejaculate during copulation. In ffSDs, the target gene is essential for female fertility but has no effect on male fitness.

The model assumes a panmictic population with discrete, non-overlapping generations. Each generation is composed of four stages: 1) eggs, 2) larvae (or juveniles), 3) virgin adults (females and males), and 4) mothers (i.e., inseminated females) which produce the next generation eggs (Fig. 2). A single release of transgenic males carrying each drive takes place at generation zero. Drives efficiency, intended effects, and unintended fitness costs, among other intrinsic drives and population features, are adjustable (see Tables 1 and 2 for a complete description of main model parameters).

**Fig. 2.**
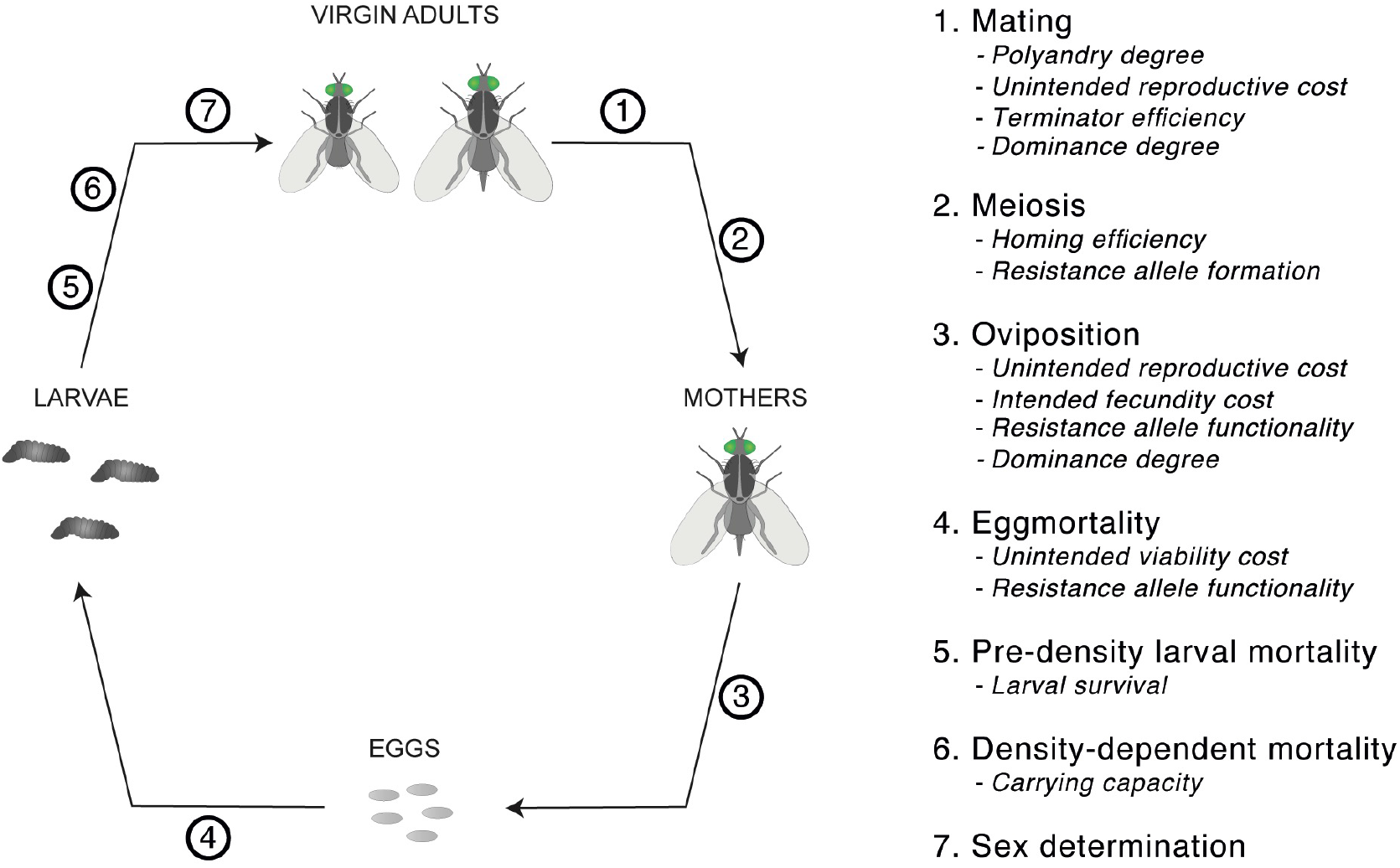
Model stages and steps across a generation. Major invoked parameters are shown per step. 1) Female and male virgin adults mate randomly giving rise to mothers, i.e., inseminated females. Females may mate more than once (*Polyandry degree*). Male mating success of drive-carrying males is reduced by unintended drive effects (*Unintended reproductive cost*), and terminator males may kill the female during mating (*Terminator efficiency* and *Dominance degree*). 2) For simplicity, meiosis is only modeled for surviving mothers and mating males (*Homing efficiency* and *Resistance allele formation*). 3) Each mother produces a clutch of eggs where the expected number of eggs may be reduced for drive-carrying mothers (*Unintended reproductive cost, Intended fecundity cost, Resistance allele functionality*, and *Dominance degree*). 4) Eggs have a probability of dying before reaching the larval stage, which may be higher for drive-carrying eggs (*Unintended viability cost* and *Resistance allele functionality*). 5) Larvae have a probability of dying right after hatching (*Larval survival*). 6) Then, larvae are subject to density-dependent mortality (*Carrying capacity*). 7) Surviving larvae result in virgin adults, and sex is randomly determined.

**Table 1.**
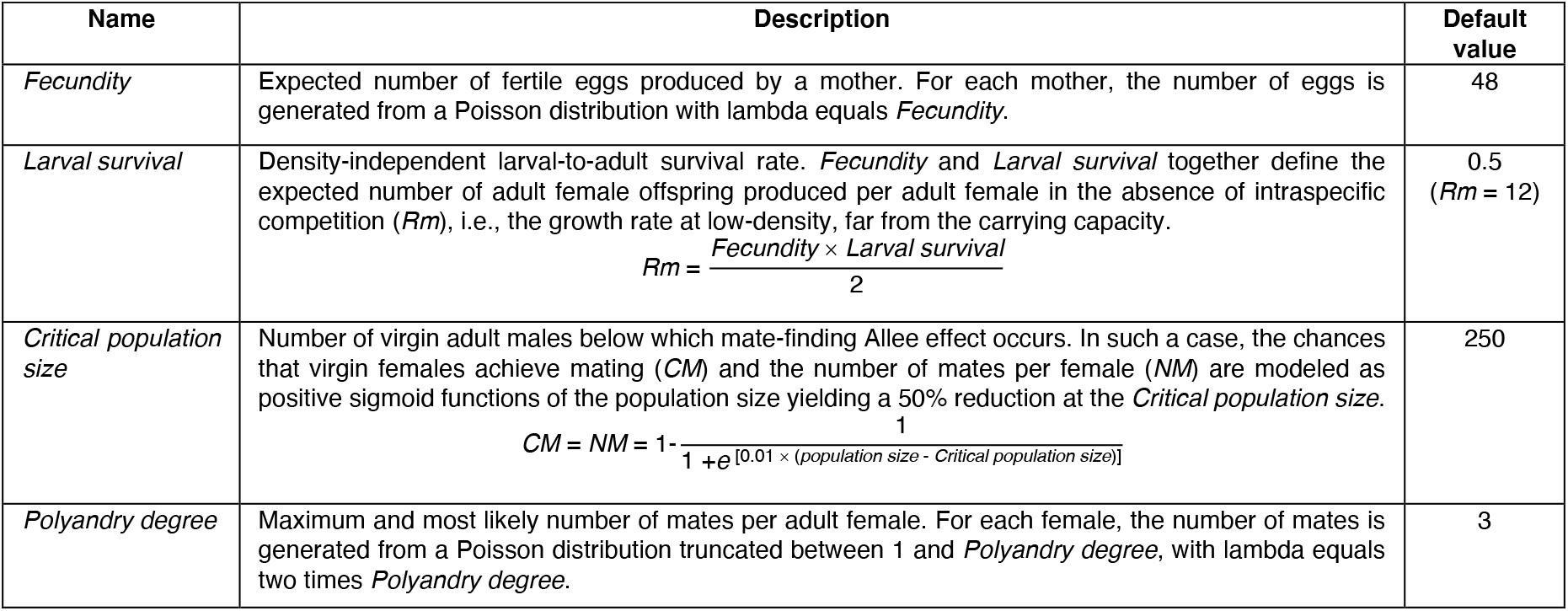
Main population parameters. A description and default value are shown for each parameter.

**Table 2.** Main system parameters. A description and the default value(s) are shown for each parameter. For the three parameters related to intended drive effects, the default value may vary between designs (ffBED, sBED, and ffSD). For the four parameters related to drive endonuclease activity and unintended fitness cost, two default values are included: one for a system with ideal drive activity [Ideal System (IS)], that has perfect conversion rate and does not impose unintended fitness costs, and the other for a system with reference drive activity [Reference System (RS)], with imperfect conversion and unintended fitness costs.

| Name | Description | Default value |  |  |
| --- | --- | --- | --- | --- |
|  |  | ffBED | sBED | ffSD |
| *Intended fecundity cost* | Relative reduction of mother's fertility (total number of fertile eggs a mother is expected to produce) caused by drive system activity. In BED designs, this reduction is experienced by mothers homozygous for both drive system components (AABB mothers, whereas A and B are each of the gene drive components). In SD designs, it is experienced by mothers homozygous for the drive (CC, whereas C is the gene drive). If *Dominance degree* is higher than 0 (see below), this cost will also be experienced, at least partially, by non-homozygous mothers carrying at least one allele of each system component in BED designs (AABb, AaBB, or AaBb mothers) or by drive-carrying heterozygous mothers in SD designs (Cc mothers) (lower case letters designate native alleles). | 1 | 0 | 1 |
| *Terminator efficiency* | Probability that a female that copulates with an AABB terminator is killed following mating due to the action of the drive system. It only applies to BED designs. If *Dominance degree* is higher than 0 (see below), *Terminator efficiency* will also apply, at least partially, for non-AABB terminators (AABb, AaBB, or AaBb males). | 0 | 0.95 | 0 |
| *Dominance degree* | Proportion of the drive system effect (*Terminator efficiency* or *Intended fecundity cost*) experienced by double heterozygotes (AaBb) carrying one allele of each drive in BED designs or by drive-carrying heterozygotes (Cc) in SD designs. Drive-carrying homozygotes experience full intended effect (see above). In BED designs, individuals homozygous for one drive but heterozygous for the other will experience an intermediate effect between the effect experienced by AaBb and AABB individuals. In SD designs, this parameter also affects the expected fecundity of heterozygous mothers carrying a resistance allele (see *Resistance allele functionality* below). | 0.6 |  | 0.2 |
| *Unintended reproductive cost* | Relative reduction of female fertility and male mating success due to unintended drive activity. For drive-carrying mothers, it is the reduction of the expected number of fertile eggs. For drive-carrying males, it is the reduction in mating success. These costs are dominant (i.e., mothers and males carrying one and two drive alleles experience the same reduction) and in BED designs are imposed by each drive separately. | 0 (IS) or 0.025 (RF) |  |  |
| *Unintended viability cost* | Probability that a drive-carrying egg perishes due to unintended drive activity. This cost is dominant (i.e., eggs carrying one and two drive alleles experience the same viability reduction) and, in BED designs, is imposed by each drive separately. | 0 (IS) or 0.025 (RS) |  |  |
| *Homing efficiency* | Probability that a drive converts the wild target locus into a drive allele during gametogenesis in drive-carrying heterozygotes. | 1 (IS) or 0.9 (RS) |  |  |
| *Resistance allele formation* | Probability that a drive converts the wild target locus into a cleavage-resistant allele (resistance allele) during gametogenesis in drive-carrying heterozygotes. | 0 (IS) or 0.05 (RS) |  |  |
| *Resistance allele functionality* | Relative functionality of resistance alleles. Ranging from 0 to 1, in BED designs (where target genes are essential for development), it is the probability that viable eggs homozygous for resistant alleles of any drive develop into a larva. In SD designs, where the target gene is essential for female fertility, it is the fraction of *Fecundity* preserved in mothers homozygous for resistant alleles. To be conservative in BED designs, the deleterious effect of resistance alleles was assumed to be recessive (i.e., resistant-wt-heterozygotes, resistant-drive-heterozygotes, and wt-homozygotes are equally functional for the target genes). In ffSD, the expected fecundity of resistant-wt-heterozygous females (or resistant-drive-heterozygous females) is intermediate between resistance-homozygous females (or drive-homozygous females, respectively) and cc females, whereas *Dominance degree* determines linearly how close fecundity of resistant-wt-heterozygotes (or resistant-drive-heterozygotes) is to resistance-homozygotes' (or drive-homozygotes', respectively). | 0 |  |  |

Inspired by the performance of previous gene drive systems in flies and mosquitoes ^7,10,18,20,26,29,30^, we choose realistic—yet optimistic—parameters values as default (Tables 1 and 2). For each modeled design, two classes of drive activity were considered: ideal and reference. In ideal systems, drive conversion rate (i.e., homing efficiency) is 100% (which implies no resistance allele formation) and drive(s) activity imposes neither reproductive nor viability fitness cost. In reference systems, homing efficiency is 90%, resistance allele formation rate is 5%, unintended reproductive cost (imposed on male mating success and female fecundity by each drive separately) is 2.5%, and unintended viability cost (imposed on eggs also by each drive separately) is also 2.5%. As both *Unintended reproductive cost* and *Unintended viability cost* were kept equal in all the performed simulations, we refer to these parameters together as *Unintended fitness costs*.

## Results

To test the performance of our model we first set our parameters to their defaults (ideal or reference) and then explored parameter space for one or two parameters at a time. For every considered scenario, we performed 100 simulations, each of which comprised up to 36 or 60 generations starting from 50,000 virgin adults and 250 adult transgenic males released per drive (0.5% of the population carrying capacity) in BED designs or 500 in SD designs. In all cases, we recorded the adult population size and allele frequencies (drives, wt, and resistance alleles) in every generation, and the rate of elimination within 36 generations, which we termed ‘fast elimination rate’. The number of replicate simulations was set to 100 since, at least with default parameters, the dispersion (difference between the minimum and maximum) of the final adult population size always stabilizes within 50 simulations.

### Homing efficiency and unintended drive effects

Fig. 3 summarizes the performance of the three gene drive designs with ideal and reference drive activity. Using the ideal drive parameters, we observed a fast and complete elimination rate for all three designs. However, using the reference drive parameters (i.e. assuming imperfect drive activity), performance differed between designs. Overall, ffBED showed the worst performance. It was not able to eliminate the target population in any simulation, although it reduced the population by ∼50% in 17 generations. Furthermore, irrespective of how low the drive costs on fitness were, the population was only eliminated under perfect homing efficiency (Fig. 3, upper panels). In contrast, sBED showed great suppression potential, tolerating a wide range of *Homing efficiency* and *Unintended fitness costs* (Fig. 3, middle panels). Therefore, further exploration of alternative scenarios and parameter space is only shown for sBED alone or together with ffSD for comparison purposes.

**Fig. 3.**
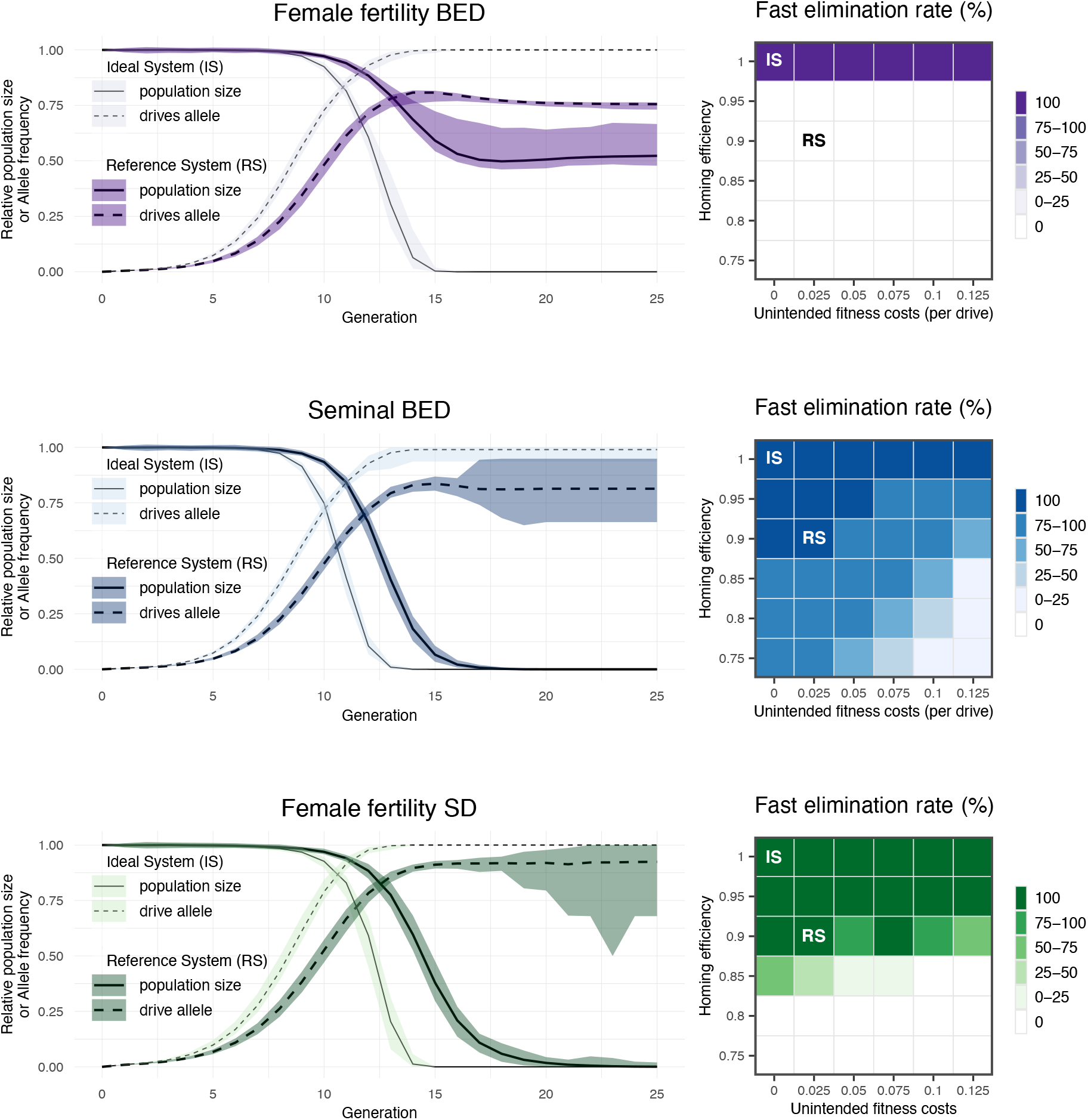
Suppression potential of ffBED (upper panel), sBED (middle panel), and ffSD (lower panel). Left: population size (relative to the initial population size) and drive(s) allele frequency are shown across the first 25 generations for each design using ideal and reference drive parameters. Lines represent the 100-simulations mean value while the colored envelopes stand for the maximum and minimum values. For BED designs, frequencies of the two drives were averaged. Right: fast elimination rate (the percentage of simulations where the population was eliminated within 36 generations) is shown for a combined space of *Homing efficiency* and *Unintended fitness costs* (*Unintended reproductive cost* and *Unintended viability cost*). *Resistance allele formation* was assumed to be half the complement of *Homing efficiency* (e.g., if *Homing efficiency* is 0.8, *Resistance allele formation* is 0.1).

In general, sBED was superior to ffSD when comparing the fast elimination rate (within the first 36 generations) across a range of *Homing efficiency* and *Unintended fitness costs* (Fig. 3, right panels). With reference drive parameters, for instance, though both designs always eliminated the population in less than 36 generations, it took ffSD on average 5.5 extra generations. sBED also succeeded in suppressing the population under conservative levels of *Homing efficiency* and *Unintended fitness costs*. For instance, when efficiency was reduced to 0.8 and *Unintended fitness costs* raised to 0.05, the intervention resulted in a population collapse in 82% of the cases in less than 24 generations. Under the same conditions, ffSD did not result in the population collapsing before generation 36 in any simulation. However, when *Homing efficiency* and *Unintended fitness costs* were set at 0.95 and 0.1, respectively (or very similar values), the fast elimination rate was slightly higher for ffSD (100%) than for sBED (99%).

### System dominance

In ffSD designs, the target should be largely haplosufficient so heterozygotes do not suffer the intended fitness cost imposed by the drive, allowing the invasion of the drive in the population. Since such a target gene may be difficult to find in some species, this requirement is an important drawback of ffSD designs. In terms of our model, haplosufficiency of the target corresponds to low values of the *Dominance degree* parameter. While the *Terminator efficiency* parameter (in sBED designs) or the *Intended fecundity cost* parameter (in ffSD designs) determines the system intended effect expected for drive-carrying homozygotes, the *Dominance degree* parameter determines the extent to which this intended effect manifests in drive-carrying heterozygotes (Table 2). In sBED designs, the default value of *Dominance degree* was 0.6, which means that adult males carrying one copy of each drive exhibit, relative to double homozygous drive-carrying males, 60% of the ability to kill females during mating. In ffSD designs, the default value was 0.2, which means that heterozygous mothers experience 20% of the fecundity reduction that is experienced by drive-homozygous mothers.

Therefore, to analyze how sBED may deal with more dominant or recessive expression systems and how ffSD may deal with more dominant or recessive target phenotypes, we further explored the performance of both designs with ideal and reference drive parameters at *Dominance degree* values of 0, 0.33, 0.67, and 1. Our results reveal that the effect of *Dominance degree* on suppression capability is less for sBED than for ffSD designs. Indeed, for sBED, *Dominance degree* did not notably affect the performance using ideal drive parameters (Fig. 4, upper panel). Even with imperfect drive activity (reference drive parameters), the sBED potential for fast suppression, relative to ffSD, persisted across a wider range of *Dominance degree* values. While the reference sBED showed fast and efficient population suppression except under very low *Dominance degree*, the reference ffSD did so for a narrower window of low *Dominance degree* values (Fig. 4).

**Fig. 4.**
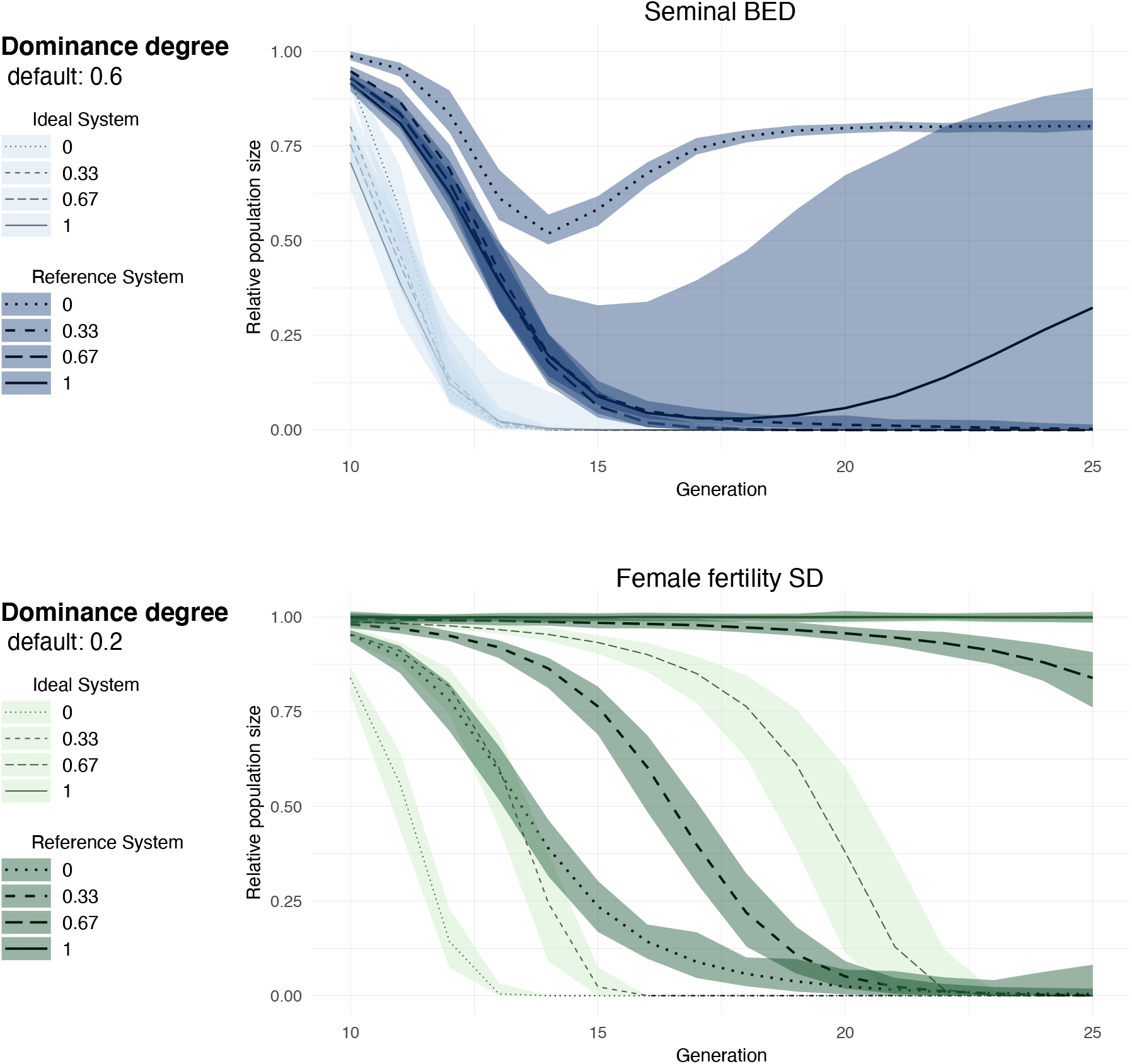
Impact of *Dominance degree*. Relative population size from generation 10 to 25 is shown with ideal and reference drive parameters for sBED (upper panel) and ffSD (lower panel) using in each case different values of *Dominance degree*. Lines represent the 100-simulations mean value while the colored envelopes stand for the maximum and minimum values. The graphs using the *Dominance degree* default values are not shown here as they are already shown in Fig. 3.

### Low-density growth rate

We refer to low-density growth rate (or *Rm*) as the expected number of adult female offspring produced per adult female in the absence of intraspecific competence. Higher *Rm* values indicate the production of greater offspring numbers per female, allowing the population to quickly recover from the deleterious effects of the drive system. Therefore, the fast elimination rate is expected to decrease with *Rm*. As previously reported, one limitation of ffSD is that it hardly eliminates populations with high *Rm* ^12^.

In our model, *Rm* can be adjusted with the *Fecundity* parameter (Table 1). Therefore, to evaluate the impact of *Rm* we explored the *Fecundity* parameter space. Our result supports the previous finding on ffSD and also reveals that sBED may deal better with high-*Rm* populations (Fig. 5). While with ideal drive parameters both designs could in all cases eliminate populations with *Rm* of 24 in less than 36 generations, with reference drive parameters the fast elimination rate was remarkably higher for sBED (100%) than for ffSD (12%).

**Fig. 5.**
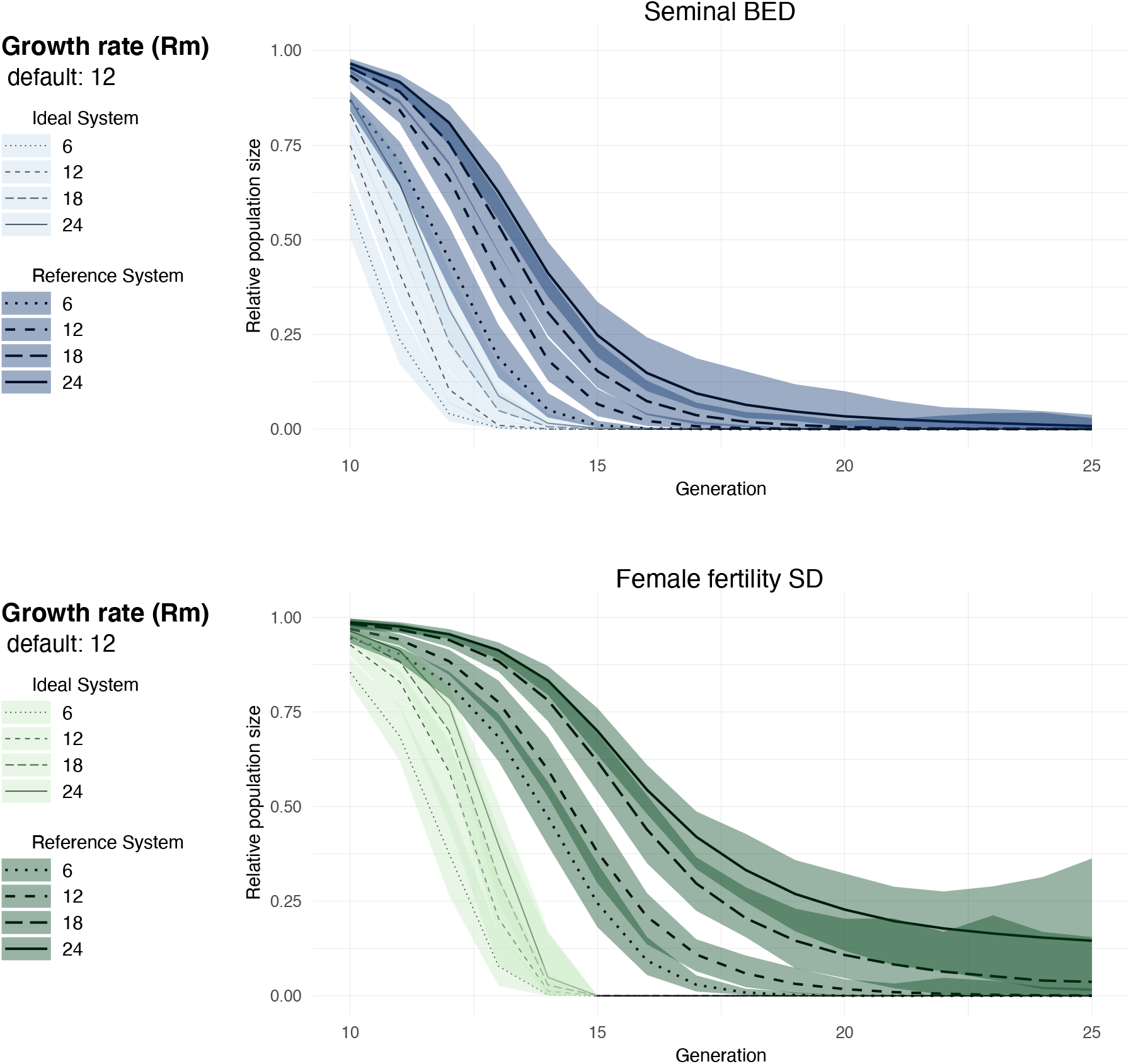
Impact of low-density growth rate. Relative population size from generation 10 to 25 is shown with ideal and reference drive parameters for sBED (upper panel) and ffSD (lower panel) using in each case different values of *Rm. Rm* was adjusted with the *Fecundity* parameter. Lines represent the 100-simulations mean value while the colored envelopes stand for the maximum and minimum values.

### Resistance allele fitness

For homing to occur in CRISPR-Cas9-mediated gene drives, the double-stranded breaks induced by Cas9 nuclease must be resolved by the homology-directed repair pathway (HDR), which predominates specifically during early meiosis ^31^. When the breaks are produced and resolved before or after the window optimal for HDR, nonhomologous end joining (NHEJ) prevails and the sequence of the target site is often changed so it is no longer recognized by the drive (e.g., ^24^). Frequently, NHEJ results in cleavage-resistant alleles (so-called resistance alleles) with disrupted gene function (also known as nonfunctional or r2 resistance alleles) that tend not to significantly affect the propagation of the drive allele since they will be selected out of the population ^13^. Alternatively, if the resistance allele preserves the function of the target, it will possess an adaptive advantage over the drive allele and thus represents a great obstacle to the spread of the drive. The formation rate of these functional resistance alleles (also known as r1 resistance alleles) is rare and can be further reduced, for instance, by using multiple gRNAs ^13^ or a conserved target gene that cannot tolerate mutations ^18^.

In our models, resistance alleles were considered nonfunctional (r2) by default. However, given that r1 resistance alleles can restrict gene drive propagation ^32^, we also assessed how our design is affected by the level of fitness reduction imposed by resistance alleles. To do so, we explored the performance of the systems assuming different degrees of *Resistance allele functionality* (see the parameter description in Table 2). Our results show that with reference drive parameters (which admits the formation of resistance alleles) sBED is more tolerant than ffSD to functional resistant alleles. While ffSD did not even tolerate a low *Resistance allele functionality* of 10%, sBED showed a fast elimination rate of 85% with as high as 40% functional resistance alleles (Fig. 6).

**Fig. 6.**
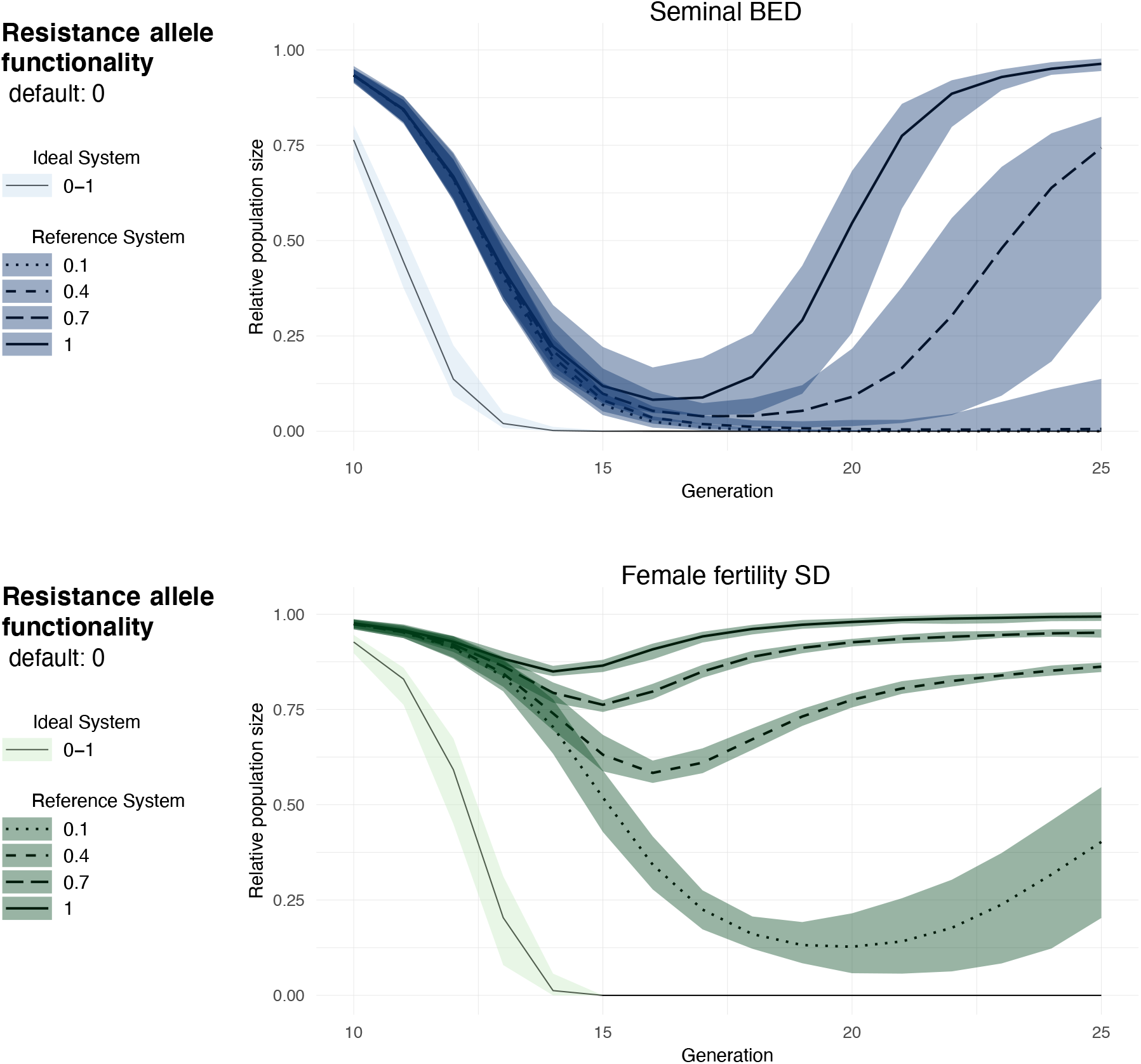
Impact of *Resistance allele functionality*. Relative population size from generation 10 to 25 is shown with ideal and reference drive parameters for sBED (upper panel) and ffSD (lower panel) using in each case different values of *Resistance allele functionality*. Lines represent the 100-simulations mean value while the colored envelopes stand for the maximum and minimum values. For reference systems, the graphs using the *Resistance allele functionality* default value are not shown here as it is already shown in Fig. 3.

### Polyandry and terminator efficiency

By default, in our model, *Polyandry degree* was set to 3, which means that each adult female copulates with one, two, or, more likely, three males (Table 1). By reducing *Polyandry degree* in sBED designs with ideal drive parameters, the fast elimination rate was not affected, yet elimination was slightly delayed (Fig. 7). In reference sBED designs, changing *Polyandry degree* from 3 to 2 resulted in a severe reduction of the fast elimination rate, from 100% to 1%. Nonetheless, in 78% of the cases, the population was eliminated between generations 36 and 60 (we did not simulate more generations). In the cases where the population was not eliminated by generation 60, it remained highly suppressed at low size (Fig. 7). When *Polyandry degree* was reduced to 1, the fast elimination rate fell to 0%, which indicates that sBED interventions may not be able to completely suppress populations with monogamous females. However, population size remained reduced at ∼53% from generation 16 (Fig. 7), resembling the output of ffBED (Fig. 3). In contrast, *Polyandry degree* did not affect the intervention outcome in ideal or reference ffSD designs. This is not surprising because ffBED affects female fertility irrespective of the number of mates.

**Fig. 7.**
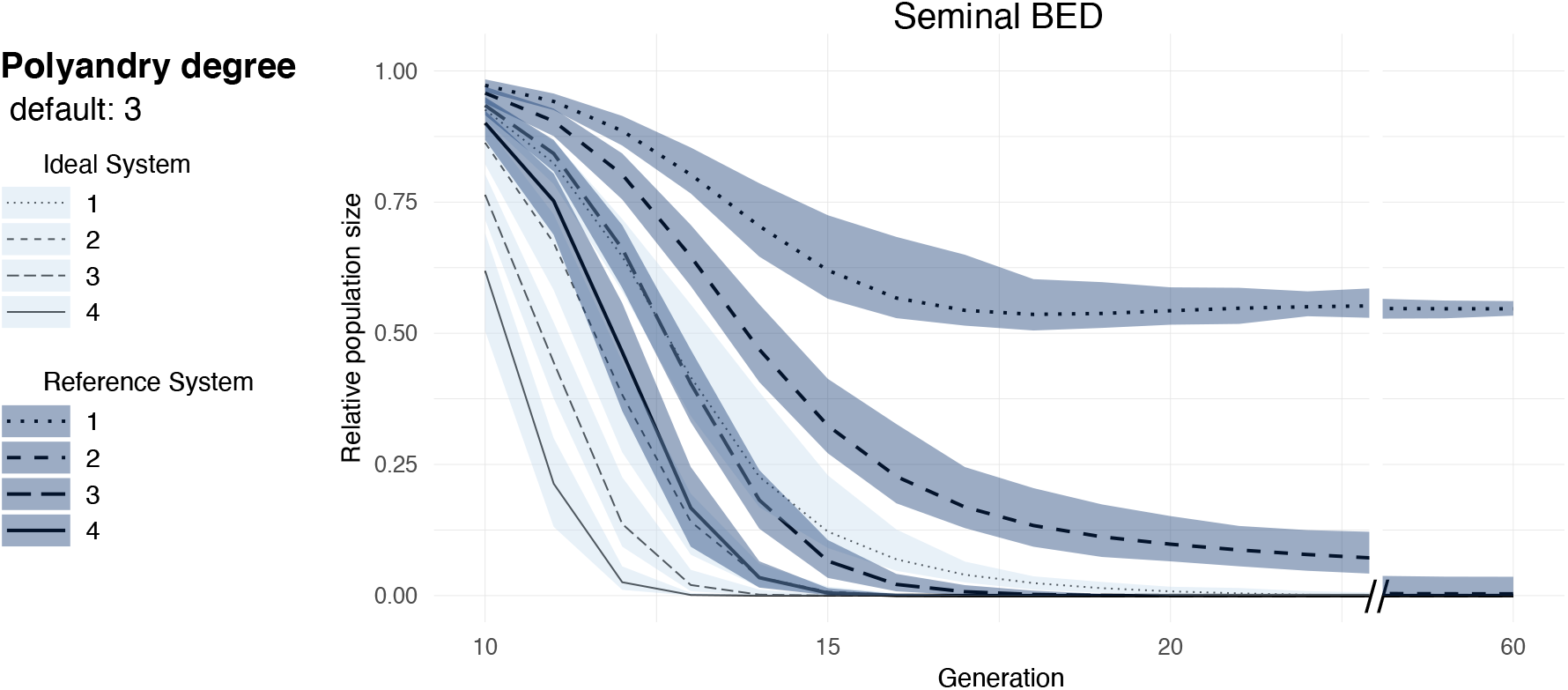
Impact of polyandry. Relative population size from generation 10 to 60 is shown with ideal and reference drive parameters for sBED (upper panel) and ffSD (lower panel) using in each case different values of *Polyandry degree*. Lines represent the 100-simulations mean value while the colored envelopes stand for the maximum and minimum values.

It can be expected that highly efficient transferred seminal toxins, whose action is modeled through the *Terminator efficiency* parameter (Table 2), may compensate for the effect of reduced polyandry. To evaluate this possibility, we estimated the fast elimination rate of the sBED with reference drive parameters under different combinations of *Terminator efficiency* and *Polyandry degree* (Fig. S1). We also repeated the simulations using different combinations of *Critical population size*, which determines the Allee effect range (Table 1), and varying the number of released males per drive. Our results showed a poor compensation capacity of *Terminator efficiency* on the effect of *Polyandry degree*. Overall, a high fast elimination rate was only achieved when *Terminator efficiency* was 0.8 (or higher) and *Polyandry degree* was 3 (or higher). Furthermore, neither *Critical population size* nor the number of released males per drive notably affected the simulation outcome.

## Discussion

Our results reveal that the use of BEDs to spread dominant or semidominant toxins has the potential for effective population suppression. Particularly, we show that the sBED design may be highly efficient and work relatively fast against polyandrous target populations. We identified, however, three key elements that may affect the feasibility of this approach: 1) the challenge of generating effective seminal bioinsecticides, 2) the requirement that target females be polyandrous, and 3) the lack of panmixia in the target population.

The development of a sBED implies the design and generation of seminal toxins that do not affect the mating success of the male that produces it but kill or sterilize the target female. Currently, many effective bioinsecticides are bacteria-derived protoxin proteins known to be proteolytically activated upon ingestion by the insect midgut lumen proteases ^33^. Interestingly, the female reproductive tract resembles the insect midgut as both tracts have a rich repertoire of secreted proteases, especially serine proteases. In the insect uterus, these proteases are responsible for processing male-derived peptides, degrading the mating plug, and protecting the female from microbes (e.g., ^34–41^). Though no protoxin has ever been evaluated inside the female reproductive tract, some insecticidal proteins may exhibit similar toxic effects therein. Also, given the cellular and physiological differences between the male accessory glands, where most seminal proteins are produced, and the uterus, it might be possible to design insecticidal proteins that will be activated solely within female tissues. Given the recent advances in knowledge of biological control agents ^42–45^, the development of in silico tools to predict and scan molecular targets for pest control ^46^, and the advances in computational protein design ^47–50^, the engineering of seminal toxins may be attainable in the short term. However, the evolution of resistance to the toxins in the target population may always restrict insecticide-based approaches (reviewed in ^51^).

Concerning polyandry in our model, the success of the sBED intervention requires that most females mate with more than one male before oviposition (Fig. 7) to allow each terminator male the opportunity to kill (or sterilize), on average, more than one female. Although multiple paternity is common among insects, the level of polyandry is highly variable between species ^52,53^, with many species rarely remating (e.g., ^54^), therefore reducing the range of targets for which this control strategy would be effective. Nonetheless, our model showed that with ideal drive parameters, which implies that terminator males do not undergo mating success reduction, even a monandrous population can be rapidly eliminated (Fig. 7). This suggests that low polyandrous populations can be efficiently suppressed as long as drive-carrying males are highly successful in achieving mating or drives’ frequencies are kept high by releasing drive-carrying males in multiple bouts across several generations.

Mathematical modeling has shown that deviations from random mating may allow the clustering of free-drive individuals which, consequently, may thwart gene-drive-mediated extinction ^12,55^. Population structure may have a particularly strong impact on the outcome of BEDs because they require that both drive components spread evenly across the population. For example, if one of the drive components is introduced in a given subpopulation several generations before the other, without imposing the intended fitness cost, it will probably spread rapidly. However, the second component will not have the chance of spreading alone because by the time it arrives in the subpopulation too many individuals will already carry the first component. Therefore the second component, and consequently the BED system, will not be able to fully invade that particular subpopulation. This scenario would potentially produce divergence among subpopulations in which different drives establish first because hybrid males will be terminators, incapable of producing any offspring. More complex simulations are necessary to help predict the outcome of BEDs in structured populations and how repeated releases of transgenic males may enhance BEDs’ propagation.

The concept we present combines three biological tools that are versatile and have been proven to act in a wide range of taxa: artificial CRISPR-Cas9 nucleases ^56^, artificial gene expression systems such as GAL4-UAS ^57^, and insecticidal proteins such as Bt-toxins or spider venom peptides ^58^. Therefore, transferring a BED design from one species to another would likely not require major adjustments in protocols. In addition, the development of insecticidal seminal proteins will provide a significant improvement to the Sterile Insect Technique, one common approach used to control insect pest populations ^59^. The strategy consists of regular releases of sterile males unable to produce viable offspring so the target population is severely reduced or eliminated. However, female remating with wt males restricts the efficacy of this approach. Furthermore, females mated to sterile males may still attempt oviposition, which could lead to crop damage. But if instead, released males were not only sterile but also delivered seminal toxins to females during mating, target females could be killed right after copulation enhancing the efficacy of the control.

There is an ever-increasing need for ecologically conscious control of pest species that have a negative impact on health, agriculture, or the environment. Although future research on seminal protein engineering is warranted, our modeling indicates that BED designs might be a highly efficient strategy for biological control.

## Methods

### Model steps

Every generation, the number of mates for each adult virgin female is generated from a Poisson distribution truncated between 1 and *Polyandry degree*, with lambda equals two times *Polyandry degree*. Therefore, the *Polyandry degree* parameter defines the maximum and most likely number of mates per adult female. After these numbers are settled, male mates of each female are randomly sampled with replacement so a given male may mate with different females.

In sBED designs, if an adult male carrying both the transactivator and effector constructs (i.e., a terminator) successfully mates, the female then has a probability of dying after mating. This probability equals the *Terminator efficiency* if the terminator male is homozygous for both drives. However, an unintended dominant reproductive cost, which is adjusted for each drive, reduces the terminators’ chances of successfully mating (see *Unintended reproductive cost* in Table 2). The *Dominance degree* parameter determines the fraction of the *Terminator efficiency* exhibited by heterozygous terminator males (see Table 2). Each surviving female, which is assumed to carry equal amounts of sperm from all her mates, will oviposit eggs. For simplicity, meiosis is only modeled for mothers and mating males. Both the probability of homing conversion and formation of cleavage-resistant alleles (so-called resistance alleles) are adjustable and restricted to the meiosis stage of drive-wt heterozygotes (see *Homing efficiency* and *Resistance allele formation* in Table 2).

Giving rise to the next generation, each mother produces a clutch of eggs, whose size is generated from a Poisson distribution with lambda equals *Fecundity*. However, according to the *Unintended reproductive cost* parameter, the fecundity of drive-carrying mothers is reduced by each drive separately. In ffBED designs, the fecundity of mothers that are homozygous for both drives (i.e., AABB mothers, whereas A and B are each of the gene drive loci) is further reduced according to the *Intended fecundity cost* parameter. The same occurs to homozygous drive-carrying mothers in ffSD designs (i.e., CC mothers, whereas C is the gene drive locus). The *Dominance degree* parameter determines the fraction of the intended fecundity cost that affects heterozygotes.

Eggs’ genotypes are stochastically generated by sampling with replacement the haplotypes of the mother’s gametes and the sperm she stores. According to the *Unintended viability cost* parameter, which defines a dominant viability cost imposed by each drive, drive-carrying eggs may have an added probability of dying before reaching the larval stage. Surviving eggs develop into larvae which are subject to density-independent and density-dependent mortality according to the Beverton-Holt model ^60^. Each surviving larva results in a virgin adult female or male with equal probability.

### Simulation process

At generation zero, the population size is in equilibrium at its carrying capacity with an equal number of females and males. Released males are homozygous (AAbb and aaBB) in BED designs while heterozygous (Cc) in SD designs, given that large amounts of homozygotes may be difficult to produce. At the virgin adult stage of every generation, if the number of males falls below a critical number (see *Critical population size* in Table 1), the population experiences a mate-finding Allee effect ^61^, making mating opportunities less likely. Specifically, we modeled both the probability that virgin females achieve mating (and become mothers) and the number of mates per mother as positive sigmoid functions of the population size (for more details see Table 1). The population simulation ends (i.e. the population collapses) when no female becomes a mother, no mother produces viable eggs, or no larva survives to become an adult virgin female.

We created a tool in R ^62^, binXdrives, to run the simulations. The tool is available upon request and will be available at https://github.com/santiagorevale/binXdrives. See software versions in Table S1.

## Code and data availability

The source code of binXdrives and all simulation data generated in support of this research is available upon request and will be available at https://github.com/santiagorevale/binXdrives.

## Acknowledgments

We thank Prof. Fred Gould and members of his lab for assistance during the initial stages of model designing. We also thank Silvina Belliard for comments that helped in different stages of the present study. This work was supported by The National Scientific and Technical Research Council (Argentina). Computational procedures of this research were supported by the Wellcome Trust Core Award Grant Number 203141/Z/16/Z with additional support from the NIHR Oxford BRC. The views expressed are those of the authors and not necessarily those of the National Health Service, the NIHR, or the Department of Health.

## Authors contributions

JH conceived the original idea, designed the model, and took the lead in writing the manuscript. JH and SR wrote the code. SR optimized the code and run the simulations. LMM was involved in planning the work and supervised the project. JH and LMM wrote the manuscript. All authors discussed the results, and read and approved the final manuscript.

