## Supplementary material for "Propagation of seminal toxins through binary expression gene drives can suppress polyandrous populations": Fig. S1

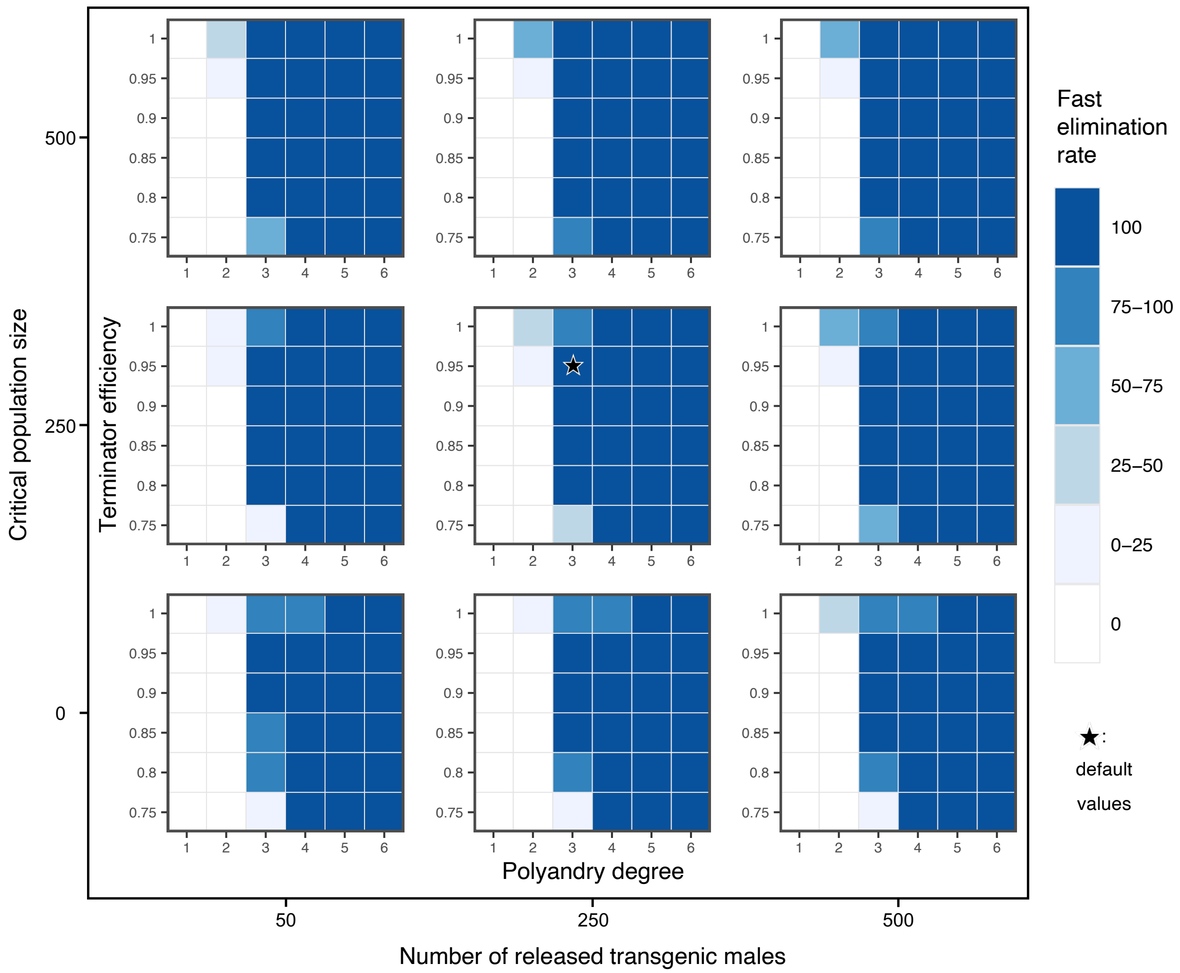


**Fig. S1 Fast elimination rate of sBED for a combined space of multiple parameters.** The percentage of simulations where the population was eliminated within 36 generations is shown for varying values of the following parameters: *Polyandry degree*, *Terminator efficiency*, *Release size* (the number of released males per drive), and *Critical population size*.

**Table S1 Versions of the implemented softwares.**

| Software | Version |
|---|---|
| R core | 4.0.0 |
| ggplot2 | 3.3.5 |
| logger | 0.2.0 |
| bettermc | 1.1.1 |
| dplyr | 1.0.7 |
| rlist | 0.4.6.1 |
| zeallot | 0.1.0 |
| stringr | 1.4.0 |
